# Genome report: Whole genome sequence and annotation of *Penstemon davidsonii*

**DOI:** 10.1101/2023.08.31.555769

**Authors:** Kate L. Ostevik, Magdy Alabady, Mengrui Zhang, Mark D. Rausher

## Abstract

*Penstemon* is the most speciose flowering plant genus endemic to North America. *Penstemon* species’ diverse morphology and adaptation to various environments have made them a valuable model system for studying evolution, but the absence of publicly available reference genomes limits possible research directions. Here we report the first reference genome assembly and annotation for *Penstemon davidsonii*. Using PacBio long-read sequencing and Hi-C scaffolding technology, we constructed a de novo reference genome of 437,568,744 bases, with a contig N50 of 40 Mb and L50 of 5. The annotation includes 18,199 gene models, and both the genome and transcriptome assembly contain over 95% complete eudicot BUSCOs. This genome assembly will serve as a valuable reference for studying the evolutionary history and genetic diversity of the *Penstemon* genus.

## Introduction

*Penstemon* (Plantaginaceae) is a genus made up of many flowing plant species (approximately 280) commonly known as the beardtongues. It is the largest plant genus endemic to North America. *Penstemon* species have diverse vegetative and floral morphology and are adapted to a wide range of environments (Nold 1999; Wolfe et al. 2006; Wilson et al. 2007; Thomson and Wilson 2008). This diversity has led to *Penstemon* emerging as a model system for the study of evolution, especially with regard to pollination syndromes (e.g., Wilson et al. 2004; Wessinger et al. 2014; Katzer et al. 2017), speciation (e.g., Straw 1955; Clark 1971; Wolfe et al. 1998; Stone et al. 2023), and repeated evolution (e.g., Wilson et al. 2007; Wessinger and Rausher 2014). However, the current lack of a publicly available reference genome for any species in the genus limits this research. Ultimately, reference genomes for species in the genus would significantly enhance our understanding of *Penstemon ‘s* biology and open up new avenues of research, such as studies of chromosomal evolution, the landscape of genetic divergence, and comparative genomics.

*Penstemon davidsonii* E. Greene is a perennial species that occurs in the alpine zone of the Sierra Nevada and Cascade Mountain ranges. It is a diploid with 8 chromosomes and a predicted genome size of 483 Mb (Broderick 2011). *Penstemon davidsonii* occurs within subgenus Dasanthera (Datwyler and Wolfe 2004; Wolfe et al. 2021), which is the outgroup to the rest of *Penstemon*. A reference genome for this clade will be particularly useful, as it will allow researchers to infer the ancestral genome for this clade and, therefore, better understand evolutionary changes that occurred within *Penstemon*.

Here we report the first genome assembly for *Penstemon davidsonii*. We use a combination of PacBio and Hi-C sequencing to assemble a de novo reference genome for this species.

## Materials and Methods

### Study organism and collection

We collected cuttings from multiple *P. davidsonii* plants along Piute Pass in the Sierra Nevada Mountain range in June 2016. These cuttings were allowed to root in water, transferred to a 1:1 mix of potting soil and pumice, and kept under controlled conditions in the Duke University greenhouse. Because *P. davidsonii* hybridizes with *P. newberryi* along Piute Pass (Clausen et al. 1940, Chabot and Billings 1972, Datwyler and Wolfe 2004, Kimball 2008), we used population genetic data from Garcia et al. 2023 and phenotypic characteristics to identify an individual with little *P. newberryi* ancestry. Of the plants collected, DNT005 appeared to be exclusively *P. davidsonii*. This individual was collected at 11598 ft (latitude: 37.2422, longitude: -118.68265) and used for subsequent sequencing.

### DNA isolation and PacBio sequencing

We finely ground flash-frozen tissue from DNT005 using a mortar and pestle and extracted DNA using a Cetyltrimethyl Ammonium Bromide (CTAB) protocol (Clarke 2009) with the following modifications. After grinding tissue, we performed three washes with a 0.35 M sorbitol buffer to remove secondary metabolites before cell lysis. We added 2% polyvinylpyrrolidone, 2% dithiothreitol, and 2% beta-mercaptoethanol to the CTAB buffer, and we incubated samples with RNAse A for 30 mins at 37°C before precipitating DNA.

We purified the DNA using AMPure beads (Beckmann Coulter, USA) and assessed quality using a Large Fragment Analysis Kit (Agilent, USA). The library was prepared using the SMRTbell Express Template Prep Kit 2.0 (Pacific Biosciences, USA) and Sequel II Binding Kit 2.0 with a peak insert size of around 18 kbp. The sequencing was performed on the Sequel II system with a movie time of 30 hours. This yielded 3,829,617 reads with a mean length of 30,922. The SMRTbell library and sequencing were done at the Georgia Genomics and Bioinformatics Core (GGBC).

### Hi-C library preparation and sequencing

We used intact leaf material and the Proximo TM Hi-C kit (Phase Genomics, USA) to generate libraries for Hi-C sequencing. In brief, nuclear DNA was crosslinked before endonuclease treatment with DpnII, and the fragmented dsDNA was biotinylated to create junctions as per Phase genomics protocols (van Berkum et al. 2010). These biotinylated fragments were purified and subjected to paired-end Illumina 2×150 sequencing yielding 141.6 million read pairs.

### Genome assembly

The genome was assembled using HiCanu v2.1 (Nurk 2020) consensus sequences and duplicate-purged using”purge_dups” (Guan et al. 2020) to reduce the assembly to haplotype as much as possible. Then we used the 3D-DNA pipeline (Dudchenko et al. 2017) with the default values for scaffolding. Specifically, we calculated the enzyme restriction sites for DpnII with respect to the draft assembly using the script generate_site_positions.py provided by the Juicer (Durand et al. 2016) toolkit. This restriction site map was used by the Juicer toolkit when aligning the Hi-C sequences against the draft assembly using bwa (Li 2013) to generate a duplicate-free list of paired alignments that were used by the 3D-DNA pipeline during scaffolding.

NextPolish (Hu et al. 2020) is a high-accuracy genome polishing tool that can incorporate both short reads and Pacbio HiFi reads. We prepared our reads for use by NextPolish as follows. The Hi-C reads were trimmed using cutadapt (Martin 2011) to remove low-quality bases and Ns at the start of reads. The HiFi reads were generated from the Pacbio subreads.bam using the Pacbio CCS14 tool. Then, we instructed the NextPolish pipeline to use bwa for the short read mapping and minimap2 in the mode designed for Pacbio CCS genomic reads for the HiFi read mapping by modifying the configuration file (the lgs_option section was removed as the raw Pacbio subreads were not being used, a hifi_option section was added and set to”-min_read_len 1k -max_depth 100” and the hifi_minimap2_options section was set to”-x asm20”). The completeness of the assembly was assessed using BUSCO v4.0.6. (Simão et al. 2015) based on evolutionarily informed expectations of gene content from the eudicot ODB10 database.

### RNA-seq and transcriptome assembly

We extracted DNA from frozen roots, leaves, and whole seedlings using a Spectrum Plant Total RNA Kit (Sigma-Aldrich, USA). Libraries were made using the KAPA Stranded mRNA-Seq Kit (Roche, Switzerland) with NEBNext Multiplex Oligos (96 Unique Dual Index Primer Pairs) for Illumina Barcodes (New England Biolabs, USA) and sequenced on an Illumina NovaSeq 6000 S2 150bp PE flow cell at the Duke Center for Genomic and Computational Biology Sequencing and Genomic Technologies Core.

The RNA-seq short reads were trimmed and quality filtered using trim_galore (Krueger 2015). All cleaned paired and unpaired reads were de novo assembled using Trinity v2.6.6 (Grabherr 2011) with default parameters (Haas et al. 2013) to generate the transcriptome. We used BUSCO in the transcriptome mode to estimate transcriptome completeness.

### Genome annotation and masking

We identified and masked sequence repeats through the following steps. First, we identified tandem repeats within the genome using Tandem Repeat Finder (TRF) v4.09 with specific parameter settings, including matching weight = 2, mismatching penalty = 5, indel penalty = 7, match probability = 0.8, indel probability = 0.1, minimum alignment score = 50, and maximum period size = 2,000 (Benson 1999). Second, we identified interspersed repeats and LTR retrotransposons using RepeatModeler v2.0.2 (Flynn et al. 2020). For the LTR transposons, we used the LTRStruct flag in RepeatModeler. Next, the identified repeat families were clustered using CD-HIT-EST v4.8.1 with 90% sequence identity and a seed size of 8 bp (Fu et al. 2012). Finally, the generated repeat library was applied to mask interspersed repeat elements in the assembly and develop repeat-annotation files using RepeatMaskerv4.1.0 (Smit et al. 2015).

To prepare for gene annotation using the MAKER pipeline (Campbell et al. 2014), we took the following steps. The cleaned paired RNA-seq reads were mapped to the genome assembly using Tophat v2.1.2 followed by gene annotation using Cufflinks v2.2.1. We predicted the tRNAs in the final assembly using tRNAscan-SE v2.0.7 with the default parameters (Chan and Lowe 2019). To identify miRNA precursor genes, we first retrieved 38,589 miRNAs precursors from the miRBase database (Griffiths-Jones et al. 2008) and clustered them separately to remove redundancies using CD-HIT-EST v4.8.1 (Fu et al. 2012), resulting in a total of 25,760 sequences. Then, these non-redundant sequences were used to identify the miRNA precursors in the *Penstemon* genome using homology-based search by BLASTN alignment tool with the default thresholds (Altschul et al. 1990).

We used the soft-masked genome generated using RepeatMasker v4.1.0 with the Repbase repeat library and all the previous annotations (i.e., TE, RNA-seq, miRNA, and tRNA) for gene prediction in the MAKER pipeline. Evidence from *ab inito* and homology-based methods were integrated to perform the final gene predictions. Specifically, the EST evidence from the RNA-seq assembly and ab initio gene predictions of the genome were used to construct the gene set using the MAKER pipeline. AUGUSTUS v3.2.3 (Stanke and Morgenstern 2005) was used for the *ab initio* gene prediction, and the BLAST alignment tool was used for homology-based gene prediction using the EST evidence in the MAKER pipeline. Exonerate v2.4.0 (Slater and Birney 2005) was used to polish and curate the BLAST alignment results.

## Results and Discussion

Our PacBio sequencing produced 117.5 Gb (3.8 million reads with a mean length of 30,922), which represents 268x average coverage of our 438 Mb final genome (Table 1). This genome size is 91% of the 483 Mb estimate based on flow cytometry (Broderick 2011).

**Table 1.** Summary statistics for the genome assembly of *Penstemon davidsonii*.

|  |  |
| --- | --- |
| Total bases | 437,568,744 |
| Contig count | 2,688 |
| Largest contig | 55,463,854 |
| N50 | 40,949,504 |
| L50 | 5 |
| N90 | 50,886 |
| L90 | 390 |
| Ns | 125 |
| GC content | 34.7% |
| BUSCO scores | C:95.3% [S:85.3%, D:10%], F:1.2%, M:3.5%, n:2326 |

BUSCO parameters are as follows: C = Complete BUSCOs; S = Complete and single-copy BUSCOs; D = Complete and duplicated BUSCOs; F = Fragmented BUSCOs; M = Missing BUSCOs; and n = Total BUSCO groups searched

Our initial assembly was 613 Mb, made up of 9,237 contigs. Duplicate purging reduced this to 438 Mb across 4,811 contigs. Then, scaffolding using the 141.6 million Hi-C read pairs further reduced the number of contigs to 2,688 (Table 1). In the final assembly, eight contigs greater than 20 mb make up more than 60% of the reference and likely represent the eight expected chromosomes. Completeness, as assessed by our BUSCO analysis, was approximately 95% (Table 1).

The assembled transcriptome consists of a total of 247,266,183 bases across 274,807 isoform”transcripts” with a 99% BUSCO score (C:99.2% [S:31.4%, D:67.8%], F:0%, M:0.8%, n:255; see Table 1 for abbreviations). These transcripts clustered into 180,352 putative genes of which 152,610 had a single isoform and 2,604 had 10 or more isoforms. Genome annotation using this transcriptome yielded slightly more than 18,000 gene models, of which slightly more than 17,000 were protein-coding genes (Table 2). We also identified 4,331 miRNAs and 901 tRNAs (Table 2).

**Table 2.** Summary statistics for the *Penstemon davidsonii* annotation.

|  |  |
| --- | --- |
| Genes | 18,199 |
| Protein-coding genes | 17,299 |
| Exons | 109,386 |
| CDSs | 108,295 |
| Expressed sequence matches | 41,594 |
| Protein matches | 417,305 |
| miRNA | 4,331 |
| tRNAs | 901 |

In our final genome assembly, we identified and masked 257,613,589 bases of repeat content, which represents 58.87% of the total sequence. This included 135,653 transposable elements, made up of mostly Ty1/Copia (14.93% of genome sequence) and Gypsy/DIRS1 (8.96% of genome sequence) LTR retroelements (Table 3). This is significantly more repeat elements than reported from reduced representation sequencing of *P. davidonsii (*Dockter et al. 2013).

**Table 3.** Summary of repetitive element content found in the *P. davidsonii* genome assembly.

| Element type | Number of elements | Percent of sequence |
| --- | --- | --- |
| SINEs | 0 | 0 |
| LINEs | 6,831 | 0.83 |
| LTR retroelements | 110,753 | 29.15 |
| DNA transposons | 18,069 | 1.74 |
| Rolling-circles | 5,414 | 0.40 |
| Small RNA | 1,225 | 0.31 |
| Satellites | 2,380 | 0.12 |
| Simple repeats | 91,183 | 1.28 |
| Low complexity | 13,875 | 0.15 |
| Unclassified | 391,416 | 24.90 |

This assembled and annotated genome of *P. davidsonii* provides a needed resource for the *Penstemon* evolutionary biology community. To date, evolutionary analysis of *Penstemon* species has relied primarily on phenotypic analysis (e.g., Castellanos et al. 2006), examination of patterns exhibited by a small number of genes (e.g., Wessinger and Rausher 2015), or of patterns discernible from reduced representation sequencing (e.g., Stone et al. 2021). As more *Penstemon* genomes come online, however, these analyses can be extended to a genomic scale, and investigators will be able to examine the evolutionary characteristics of entire *Penstemon* genomes.

Nevertheless, there is still room for improvement of the *P. davidsonii* genome. In particular, devolving a chromosomal-level assembly from these scaffolds will be necessary for determining properties such as recombination rate variation across the genome (e.g., Smukowski and Noor 2011) and addressing issues such as the evolution of structural variants (e.g., Ostevik et al. 2020) and genomic patterns of introgression (e.g., Martin and Jiggins 2017).

## Data availability

The raw sequencing data (PacBio WGS, Hi-C, & RNA-seq), transcriptome, genome assembly, and genome annotation have been deposited into the NCBI Sequence Read Archive under the accession number PRJNA1010203.

## Acknowledgments

The authors would like to thank Kari Ostevik and Kimmy Stanton for their help in the field and Ben Stone for suggestions about this project. We also thank the staff at the Duke Greenhouses for plant care and maintenance, the staff at IgenBio for Hi-C sequencing and help with scaffolding, and the Duke Center for Genomic and Computational Biology Sequencing and Genomic Technologies Core for sequencing and guidance. The authors also thank the members of the Georgia Genomics and Bioinformatics Core (GGBC) as well as the Georgia Advanced Computing Resource Center (GACRC) for their help.

